# Mitochondrial Methylation Two-Peak Profile Absent in Parkinson’s Disease Patient

**DOI:** 10.1101/197731

**Authors:** André X. C. N. Valente, Qiwen Liao, Guy Rohkin, Raquel Bouça-Machado, Leonor Correia Guedes, Joaquim J. Ferreira, Simon Ming-Yuen Lee

**Affiliations:** Center for Neuroscience and Cell Biology, University of Coimbra, Cantanhede, Portugal; Institute of Fundamental Medicine and Biology, Kazan Federal University, Kazan, Russia; Biocant - Biotechnology Innovation Center, Cantanhede, Portugal; State Key Laboratory of Quality Research in Chinese Medicine, Institute of Chinese Medical Sciences, University of Macau, Macao, China; Girihlet Inc., Oakland, California, USA; Clinical Pharmacology Unit, Instituto de Medicina Molecular, Lisbon, Portugal

## Abstract

Whole-mitochondrial genome methylation profiles were obtained for mitochondrial DNA (mtDNA) from blood samples of one sporadic Parkinson’s Disease (PD) patient and one healthy control donor, via Whole-Genome Next-Generation Bisulfite Sequencing. Methylation frequency was determined at 836 CpG sites out of the 1146 CpG sites in mtDNA (73% of the total). The control mtDNA methylation profile exhibited a two-peak frequency distribution, with most CpG sites showing either no methylation, or a methylation above 7%. Instead, the sporadic PD mtDNA methylation profile exhibited a generic bell-shaped frequency distribution. The data is suggestive of the bell-shaped methylation profile arising via a degradation towards randomness of the healthy sample two-peak methylation pattern. Overall, this finding provides a possible explanation for the repeated observation of PD-phenotypic characteristics in cybrid cell lines where solely the mtDNA originates from a PD patient. We discuss the finding in terms of the sporadic PD Hematopoietic Origin Theory (HOT).

## INTRODUCTION

**Figure 1.**
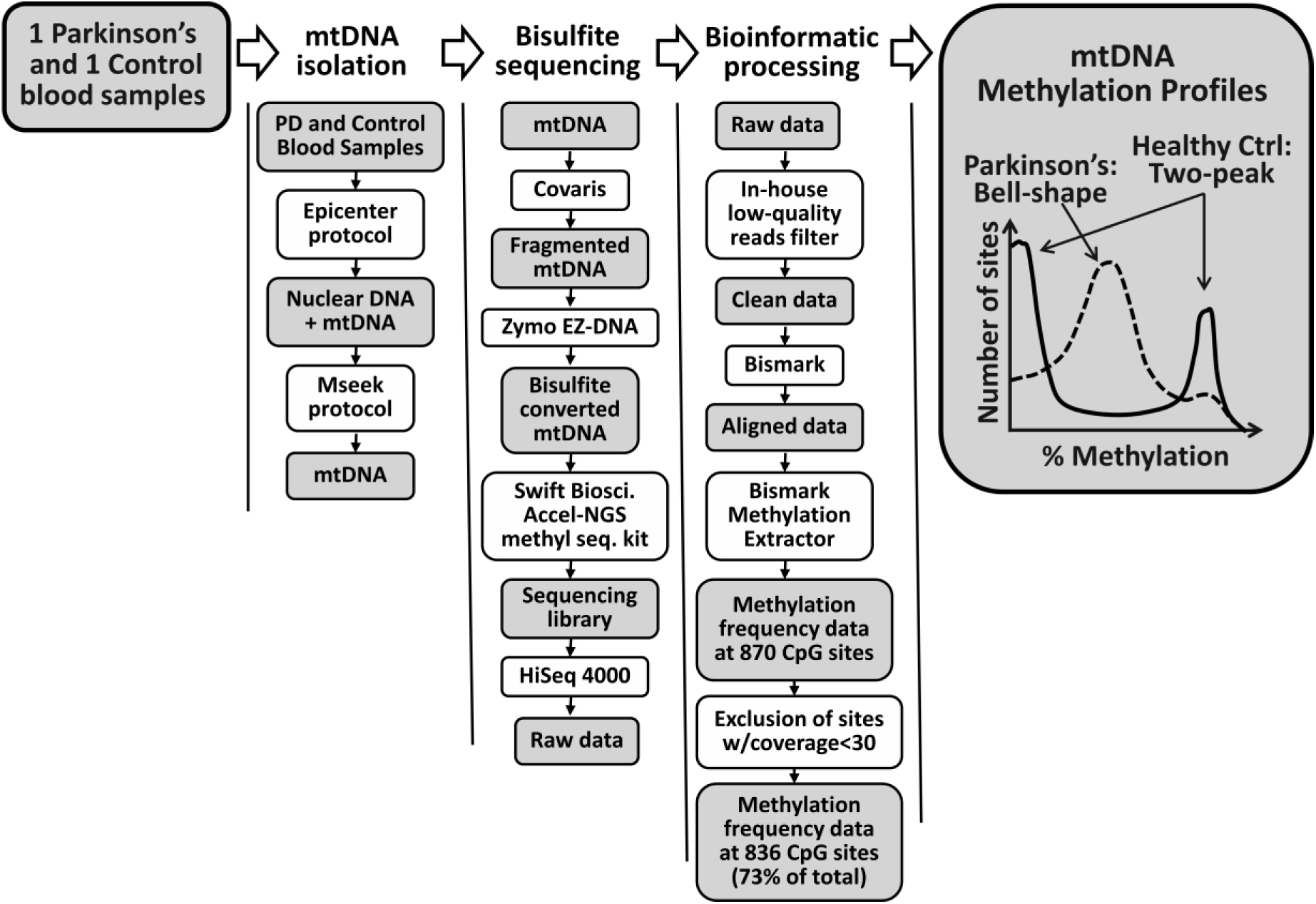
Study overview.

Parkinson’s Disease (PD) is a movement disorder characterized - and clinically diagnosed - by bradykinesia associated with rest tremor and/or rigidity (1). Existing medications and treatments only provide symptomatic relief, as there is currently no cure for PD. Approximately 10% of PD cases have a monogenic nature. Mutations in 13 genes have been associated with these Mendelian inheritable cases of PD, via familial genetic linkage studies. For non-monogenic PD, dozens of single-nucleotide polymorphisms (SNPs) risk factors have been identified based on Genome-Wide Association Studies (GWASs) (2,3,4). Likewise, a number of environmental risk factors (e.g., exposure to metals and pesticides) have been uncovered by epidemiological studies (5). However, the differential risk provided by these SNP and environmental factors is mostly very small. Thus, the roughly 90% of PD cases that are non-monogenic have an essentially unknown etiology and are clinically classified as sporadic PD. It is clear that the PD motor dysfunctions are a direct consequence of the death of the dopamine-producing neurons in the substantia nigra pars compacta region of the midbrain. However, there is no consensus on the causal chain of events between disease onset and neuronal death. In any case, the disease timeline appears to be long, as evidenced by hyposmia (6) and gastrointestinal dysfunction (7) being commonly reported as much as a decade before alterations at the motor level are noticeable. In this context, a broad spectrum of hypotheses on the origin and subsequent cascade of events in sporadic PD are still being explored. Of relevance to this article are those where mitochondria play a fundamental role.

Mitochondrial dysfunction is well-established at both the neuronal and systemic level in sporadic PD (8). However, there is no agreement on whether the observed mitochondrial alterations play a causal role in the disease, or rather are just a secondary effect of the pathological process. The standard causality argument is centered on the age-related decline in cell mitochondrial function (9,10,11,12). Key support for a causative role of mitochondria in sporadic PD comes from experiments with cytoplasmic hybrids, or ’cybrids’ (13). A cybrid is a cell created in vitro by the fusion of a neuronal cell depleted of endogenous mtDNA with an enucleated platelet. In principle, though not fully consensual, after several cybrid cell division cycles the only remaining original platelet component would be the self-perpetuating mtDNA (14). Now, various PD characteristic alterations are observed in cybrid cell lines where the precursor platelet comes from a sporadic PD patient. Therefore, cybrid experiments appear to implicate, not just mitochondria, but specifically mtDNA in the etiology of sporadic PD. On the other hand, the few mtDNA variants statistically associated with PD confer at most a very small differential disease risk and none has been unequivocally confirmed (15). Finding alterations at the epigenetic level in mtDNA from PD affected individuals would provide a possible explanation for this conundrum.

We have proposed an alternative theory on the origin of sporadic PD, which we term here the Hematopoietic Origin Theory (HOT) (3,16,17). Sporadic PD would be characterized by two phases: an induction phase, followed by a systemic propagation phase. The induction phase could occur in the hematopoietic stem-cell niche, due to an age-associated, mitochondrial-centered disruption in homeostasis. Hematopoietic cells would then acquire the PD-state at hematopoiesis. In the propagation phase, the PD-state would be transmited to other cells in the organism. We suggested that intercellular exchange of mtDNA (18,19) could be responsible for the systemic propagation of the PD-state. An imprinting of the PD-state at the mtDNA epigenetic level could prove essential in the context of such an explanation.

One well-established form of nuclear DNA epigenetic modification is methylation. Masliah et al. report a unique, concordant, pattern of nuclear DNA methylation in post-mortem frontal cortex and peripheral blood leukocyte samples from PD patients (20). Whether mtDNA can likewise be subject to methylation is currently a matter of debate. Hong et al. argue against this possibility (21). In favor of it, there is an increasing number of studies reporting the observation of mtDNA methylation, albeit at rather low levels. Bacarelli et al. report an altered methylation pattern in platelet mtDNA of cardiovascular patients (22). On PD, a recent small-scale study based on pyrosequencing by Blanch et al. reports that the D-loop region of mtDNA extracted from postmortem substantia nigra brain samples of PD patients is undermethylated by comparison with control mtDNA (23). Ultimately, we anticipate that as sequencing technologies and protocols further mature and technical issues are resolved the existence of mtDNA methylation and the validity of such studies will be conclusively settled. We mention two other open matters. First, the current research focus is on methylation at CpG sites, non-CpG methylation being mostly considered non-existent or non-biologically relevant, both for nuclear and mitochondrial DNA. This view is now being increasingly questioned (24). Second, additional mtDNA epigenetic modifications, such as 5-hmC, 5-fC and 5-caC have been identified and may yet prove to be both a common occurrence and of biological relevance (25).

Bearing the above limitations in mind, we report in here the whole-mitochondrial genome CpG methylation profiling of mtDNA purified from blood samples of one sporadic PD patient and one healthy control donor. The profiling was done via Whole-Genome Next-Generation Bisulfite Sequencing. Methylation frequency was determined at 836 CpG sites out of the 1146 CpG sites in mtDNA (73% of the total). We note that the bisulfite treatment approach we utilized does not allow differentiation between methylation proper and the related but distinct 5-hmC epigenetic modification (25). Throughout the article we shall overlook this difference and refer to the recorded epigenetic modification simply as ’methylation’.

The rest of the article is organized as follows. In *Materials and Methods* we describe the experimental procedure utilized to obtain the sporadic PD and control mtDNA methylation profiles. Proper sample purification of mtDNA is a crucial technical issue in these studies. Inadequate mtDNA isolation allows nuclear mitochondrial DNA sequence (NUMTS) regions present in nuclear DNA to confound results (26). In this regard, we utilized Mseek, a recently developed approach that has provided superior results at isolating mtDNA (27). A bisulfite treatment followed by Whole-Genome Next-Generation Sequencing approach was applied to the isolated mtDNA. In *Results*, we present the sporadic PD and control methylation profiles and our fundamental finding on their structural difference at the methylation frequency distribution level. In *Discussion*, after reviewing potential disqualifiers of the relevance of this finding, we give our perspective on how it may bear on the understanding of sporadic PD and of neurodegenerative diseases at large.

## MATERIALS AND METHODS

### Blood samples

One 400 μL PD blood sample and one 400 μL control blood sample were obtained from Biobanco-iMM, Lisbon Academic Medical Center, Lisbon, Portugal. The PD sample was from a male PD patient with no family history of PD. First symptoms of PD were detected at age 63. The sample was collected at age 73. The control sample was from a healthy female donor, with no family history of PD. The sample was collected at age 68. The study was approved by the Institute of Molecular Medicine/Santa Maria Hospital Bioethics Internal Review Board (case reference 457/15).

### Isolation of mtDNA

#### Isolating the mtDNA

Total DNA samples were obtained from each of the 400 μL blood samples via the Epicenter protocol (MC85200). Each total DNA sample was eluted in 75 μL of TE Buffer and checked for quantity and quality with a 2% Agarose E-gel (Thermo Fisher - G521802) and the Qubit dsDNA High Sensitivity assay kit (Thermo Fisher). An mtDNA sample was obtained from each total DNA sample via three successive digestions with the Exonuclease V enzyme (ExoV, New England Biolabs - M0345S). The first digestion duration was of 48h, the second and third digestions durations were of 24h each. Digestions were performed at 37 **̊**C. Between digestions, the samples were heat inactivated at 70 ̊C for 30 minutes, purified via a 1:1 wash with Ampure XP beads (Beckman Coulter) and eluted in 35 μL of PCR-grade water. Details on this Mseek mtDNA isolation protocol can be found in Jayaprakash et al. (27).

#### Confirming the absence of nuclear DNA contamination

The mtDNA samples were subjected to PCRs with the 18S and mt31 primer pairs (Table 1), utilizing OneTaq DNA Polymerase (NEB-M0480S). The 18S primer pair detects a sequence found in the nuclear genome, while the mt31 primer pair detects a sequence found in the mitochondrial genome. Reactions contained 2 μL of 10 uM of 18S or mt31 primers, 1 μL of the sample, 10 μL of OneTaq DNA Polymerase (NEB - M0480S) and 7 μL of PCR-grade water, all mixed in 0.2 mL PCR-tubes. The PCR thermocycler runs were as follows: 30 seconds at 94 **̊**C for the initial denaturation; then 34 cycles of 15 seconds at 94 **̊**C, 30 seconds at 60 ̊C and 45 seconds at 68 **̊**C; lastly 5 minutes at 68 ̊C for the final extension. Samples were then held at 4 **̊**C. The PCR products were run on a 2% Agarose E-gel (Thermo Fisher - G521802). Gel bands confirmed contamination with nuclear DNA to be insignificant in both samples (had nuclear contamination been present, an additional Exo V digestion would have been performed).

**Table 1.**
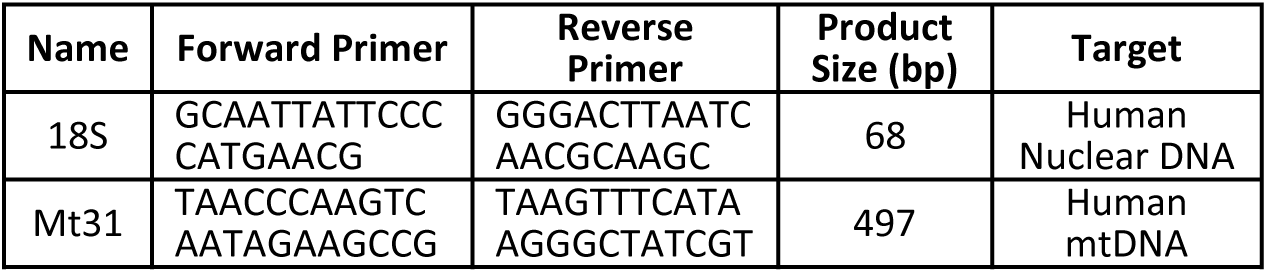
PCR primers utilized to exclude potential mtDNA sample contamination with nuclear DNA.

#### Quantifiying the isolated mtDNA

The total amount of purified mtDNA was quantified using the Qubit dsDNA HS assay kit (Thermo Fisher). The control sample yielded 12 nanograms of mtDNA and the PD sample yielded 15.6 nanograms of mtDNA.

### Whole-Mitochondrial Genome Next-Generation Bisulfite Sequencing

The starting point for this step were the 12 nanograms of control sample mtDNA and the 15.6 nanograms of sporadic PD sample mtDNA obtained from the above described mtDNA purification procedure. In brief, first the mtDNA was fragmentated to 350 bp by Covaris. Second, the EZ DNA Methylation-Gold^TM^ Kit (ZYMO RESEARCH, USA) was used for bisulfite conversion following the manufacturer’s instructions. Third, the sequencing library was constructed using the Accel-NGS^®^ Methyl-Seq kit (Swift Biosciences, USA) following the kit instruction for low input DNA. A 150 bp pair-end sequencing was then performed on a HiSeq 4000 automatic sequencing platform.

### Bioinformatic processing

The paired-end sequencing reads were filtered by an in-house C++ script with the aim of excluding low-quality and contamination reads. Clean reads were aligned with Bismark v0.17.0 (parameters: bowtie2 - n 1), with allowance for one non-bisulfite mismatch per read (28). Methylation calls were extracted with bismark_methylation_extractor. Finally, only sites with coverage of at least 30 reads in both the PD and the control samples were kept (34 sites discarded). The final result of the procedure was a dataset of methylation frequency at each of 836 CpG sites for the mtDNA purified from the sporadic PD blood sample and similarly for the mtDNA purified from the healthy control donor blood sample. In the next section, this dataset is termed the Full Dataset.

## RESULTS

### Full Dataset

The Full Dataset provides methylation frequency at each of 836 CpG sites for the mtDNA purified from the sporadic PD blood sample and similarly for the mtDNA purified from the healthy control donor blood sample. All sites have a coverage of at least 30 reads in both PD and control samples.

The healthy control sample site methylation frequency distribution (Figure 2: Full Dataset methylation histogram, black bars) is characterized by peaks at the extremes, one at the no methylation bin, the other at the methylation above 7% bin. Borrowing the terminology utilized in our mathematical work on network analysis (29), we term this a two-peak frequency distribution. In contrast, the methylation profile for the sporadic PD sample (grey bars) exhibits a bell-shaped frequency distribution. Regarding the control sample peak at the >7% bin, note that the underlying relevant fact is the presence of a significant number of sites with a frequency above 7% that is not reproduced in the sporadic PD sample. Further solidifying this difference, the average methylation frequency within the >7% bin is of 11.8% in the healthy control sample versus of 8.8% in the sporadic PD sample.

**Figure 2.**
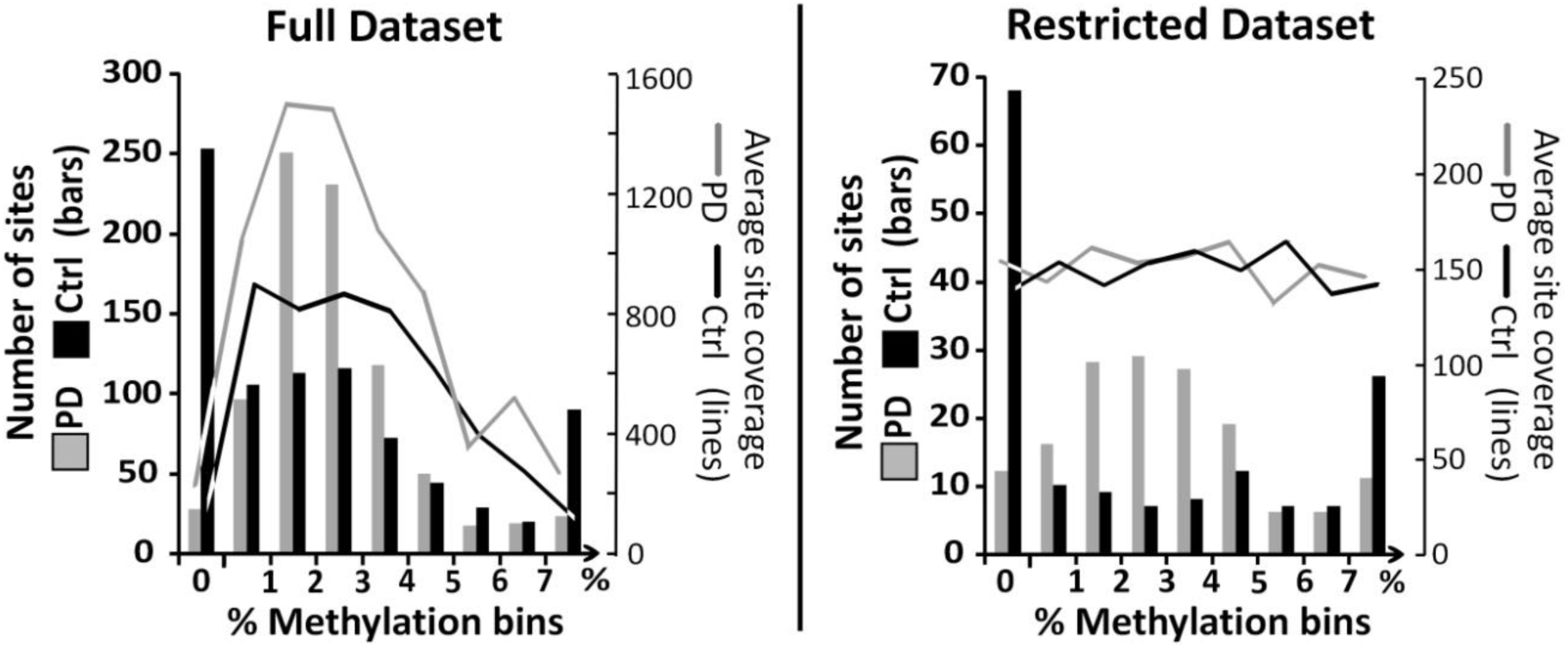
Full Dataset methylation histogram (left y-axis, bars). Minimum site coverage enforced: 30 in both samples. Total CpG sites: 836. The 0% leftmost bin contains the sites with zero methylation reads. The next bin contains the sites with methylation frequency in the (0%, 1%) range, the next in the [1%-2%) range, and so on henceforth. The rightmost bin contains all sites with a methylation frequency of 7% or above. The control mtDNA methylation profile exhibits a two-peak frequency distribution, with one peak at the no methylation bin and the other at the methylation above 7% bin. The sporadic PD mtDNA methylation profile exhibits a bell-shaped frequency distribution. **Average site coverage (right y-axis, lines).** The average coverage for the sites in each methylation bin is shown. **Restricted Dataset methylation histogram.** The Restricted Dataset is limited to CpG sites with coverage in the [105, 200] range. This coverage filter is applied separately to each sample. Total CpG sites: 154 (the coverage range was selected such as to keep an identical number of sites in the two samples). The healthy control two-peak methylation profile and the sporadic PD bell-shaped methylation profile are maintained in the Restricted Dataset. **Average site coverage.** The average site coverage is now approximately uniform, both across methylation bin and across sample.

Overall, the average methylation frequency is 2.7% in the healthy control sample and 2.5% in the sporadic PD sample, thus approximately identical. Significantly, the few sporadic PD sample >7% methylation bin sites already averaged a high 6.7% methylation frequency in the healthy control sample. In turn, the few sporadic PD sample no methylation bin sites already averaged a low 2.1% methylation frequency in the healthy control sample. This is suggestive of the ordinary bell-shaped methylation profile in the sporadic PD sample arising via a degradation towards randomness of the two-peak methylation pattern observed in the healthy control sample.

Average site sequencing coverage was 1214 reads in the sporadic PD sample versus 524 reads in the healthy control sample, a significant difference in coverage. The presence of statistical correlations between coverage and methylation frequency is clear (Figure 2: Full Dataset, average site coverage lines). In theory, the coverage differential between the two samples coupled with these correlations between coverage and methylation frequency could be the technical source of the observed distinct sample methylation profiles. To investigate this possibility, we constructed a smaller Restricted Dataset, which we discuss next.

### Restricted Dataset

The Restricted Dataset is constructed by keeping only Full Dataset CpG sites with coverage in the [105, 200] range. This coverage range filter is applied separately to each sample, so that the set of sites is now distinct in the two samples (29 sites remained in common). The coverage range was selected to yield an identical number of sites in the two samples (154 sites).

In the Restricted Dataset, average methylation frequency continues to be very close in both samples (3.2% in the healthy control sample, versus 3.0% in the sporadic PD sample). As sought, average coverage is now approximately identical in the two samples (144 in the healthy control sample versus 155 in the sporadic PD sample). The coverage is furthermore now approximately uniform across methylation frequency bin (Figure 2: Restricted Dataset, average site coverage lines). Nevertheless, the two-peak healthy control methylation profile and the bell-shaped sporadic PD methylation profile are preserved in the Restricted Dataset (Figure 2: Restricted Dataset, methylation histogram bars). Similarly, average methylation frequency in the >7% bin continues to be higher in the healthy control sample (10.9%) than in the PD sample (8.5%). In sum, the observed structurally distinct methylation profiles do not appear to be conditional on differences in sample sequencing coverage or on correlations between coverage and methylation frequency.

## DISCUSSION

A two-peak versus a generic bell-shaped structural difference between a healthy control and a sporadic PD blood mtDNA methylation profile is the fundamental finding reported in this article. Similar mean methylation levels and a tendency for the remaining PD sample high- and no-methylation sites being already so in the healthy sample are suggestive of the bell-shaped profile arising via a (partial) loss of specificity in the methylation process (see Results - Full Dataset). Next, after reviewing potential disqualifiers of the relevance of this finding, we present our viewpoint on how it may bear on the understanding of sporadic PD and of neurodegenerative diseases at large.

### Limitations

The relevance of our reported finding could be precluded three ways. First, the finding could be a technical false-positive result, the methylation profiles being in reality essentially identical in the two samples. Via the Restricted Dataset, we tried to exclude differences in coverage across sample and coverage-methylation statistical correlations as an explanation for the observed distinct profiles. However, the technologies and protocols utilized are too novel to rule out a technical false-positive due to ’unknown unknowns’. Second, the finding could be true, yet unrelated to the sporadic PD condition. We note that the two volunteers are not twins, differ in sex, are five years apart in age and could have other (clinically undiagnosed) conditions associated with them. Third, the finding could be both true and a result of the sporadic PD condition of one of the individuals, yet it could represent a secondary event, outside the causal chain of events constituting sporadic PD. Favoring a non-causative interpretation are the overall rather low levels of mtDNA methylation observed. Altogether, only the maturing of the relevant sequencing technologies and protocols together with independent analyses of additional patients will gradually settle the discussed possibilities. We presume our finding will prevail over these potential disqualifiers in the viewpoints presented next.

### On cybrids

Our finding provides a possible solution to the cybrid conundrum described in the introduction. It furthers the extensive work of PD cybrid researchers arguing for the sufficiency of mtDNA to trigger the PD-state in a cell by presenting epigenetic loss of the mtDNA two-peak methylation profile a a concrete possible trigger, in lieu of some presumed mtDNA alteration at the genetic level.

### On sporadic PD HOT (Hematopoietic Origin Theory)

Under sporadic PD HOT, a disruption in the mitochondrial dependent ROS (Reactive Oxygen Species) regulation of the hematopoietic stem-cell niche could be the most common generator of the primordial PD-state cell in an individual (17). Our observed structural alteration at the mtDNA epigenetic level in blood cells from a sporadic PD patient is thus certainly consistent with this induction phase proposed under sporadic PD HOT. Our study further bears on how the subsequent systemic propagation of the PD-state occurs. We had proposed that mtDNA itself, via its intercellular exchange, could be the vehicle for the PD-state propagation. Now we see that this propagation could specifically rely on epigenetically altered systemically disseminating mtDNA. Still unverified is the pervasiveness of mtDNA intercellular exchange. There has been renewed confirmation that mtDNA is able to move across cells under specific experimental setups, but how common such dynamics are under general physiological conditions is unknown (18,19).

### On inflammation

Our blood sample based finding reinforces the connection between sporadic PD and the hematopoietic system. However, this connection is commonly ascribed to the inflammatory process and the direct damage to cells caused by it (30). In contrast, sporadic PD HOT views inflammation, in particular chronic inflammation, as contributing to the pathology by accruing to the ’lifetime mileage’ of the hematopoietic stem cell (HSC) niche. The added stem cell work required would accelerate the eventual niche disregulation capable of generating a primordial PD-state cell. Conversely, the discovered *cumulative* PD protective effect of long-term Non-Steroidal Anti-Inflammatory Drugs (NSAIDs) intake, smoking and caffeine (31) would result from an effective ’mileage reduction’ in the HSC niche. The differential sporadic PD risk produced by certain genetic variants in the HLR region (4) would likewise follow from affecting niche mileage. Finally, the fact that PD patients are over five times more likely to be carrriers of the mutated form of GBA responsible for Gaucher’s autosomal recessive disease (32) could plausibly be due to damage caused to the HSC niche, in light of the low blood platelet levels, anemia and acumulation of the glycolipid glucocerebroside in the bone marrow, liver and spleen characteristic of Gaucher’s disease (33).

### On other neurodegenerative diseases

Neurodegenerative diseases share a variety of common features. For instance, cybrid based findings that parallel the ones discussed here for sporadic PD exist for sporadic Alzheimer’s Disease, Progressive Supranuclear Palsy, Amyotrophic Lateral Sclerosis and Huntington’s Disease (34). Thus, although our long-term research focus has been on developing a sporadic PD HOT, it is likely possible to develop related HOTs for some of these other diseases. In this regard, one issue raised by our present article finding is how does the (presumed) loss of the two-peak mtDNA methylation profile differ from one disease to another, as presumably it must.

## ACKNOWLEDGMENTS

The authors would like to thank Abhijit Sarkar (Physics, Catholic University of America) and Ravi Sachidanandam (Oncological Sciences, Icahn School of Medicine at Mt. Sinai) for manuscript criticism; Joo Heon Shin (Lieber Institute for Brain Development, Johns Hopkins University) for bioinformatic advice; Anitha Jayaprakash (Girihlet Inc.) for mtDNA isolation support; Ângela Afonso and Fabiana Rodrigues (Biobanco-iMM, Lisbon Academic Medical Center) for handling of samples; and Edson Oliveira (Neurosurgery, Santa Maria Hospital) for enabling the collaboration with Santa Maria Hospital.

## FUNDING

This work was supported by the Montepio Foundation; the Center for Neuroscience and Cell Biology (ref. PEst-C/SAU/LA0001/2013-2014); Portuguese national funds via the programs FEDER and COMPETE; the Fundação para a Ciência e Tecnologia; the Program for the Competitive Growth of Kazan Federal University; a state subsidy to Kazan Federal University in the sphere of scientific activities; the Science and Technology Development Fund of Macao SAR (ref. FDCT134/2014/A3); and the University of Macau Research Committee (ref. MYRG2016-00129-ICMS-QRCM).

